# High-throughput Single-cell Proteomics Enabled by Integrating nPOP workflow with Quantitative Hyperplexing

**DOI:** 10.64898/2026.03.05.709467

**Authors:** Kunpei Cai, Qing Zeng, Chuanxi Huang, Jing Yang, Fuchu He, Yun Yang

**Affiliations:** Department of Chemistry, College of Science, Southern University of Science and Technology, Shenzhen 518055, China; International Academy of Phronesis Medicine (Guang Dong), Guangzhou 510320, China; Guangzhou National Laboratory, Guangzhou 510005, China; Guangzhou Municipal and Guangdong Provincial Key Laboratory of Molecular Target & Clinical Pharmacology, the NMPA and State Key Laboratory of Respiratory Disease, School of Pharmaceutical Sciences, Guangzhou Medical University, Guangzhou 511436, China; State Key Laboratory of Medical Proteomics, National Center for Protein Sciences (Beijing), Research Unit of Proteomics Driven Cancer Precision Medicine (Chinese Academy of Medical Sciences), Beijing 102206, China

**Author notes:** **Corresponding Authors: Yun Yang**, **Fuchu He**.

## Abstract

Single-cell proteomics (scProteomics) has emerged as a powerful approach to dissect cellular heterogeneity and dynamic molecular mechanisms at unprecedented resolution. However, achieving high proteome coverage and quantitative accuracy while maintaining high-throughput remains a major challenge. In this study, we established a high-throughput scProteomics workflow that integrates a modified nano-proteomic sample preparation (nPOP) workflow with an IBT16 and TMT16 quantitative hyperplexing strategy. Through systematic optimization of chromatographic and mass spectrometric conditions, we established a label-free workflow for high-sensitivity and high quantification accuracy. On average, more than 3,000 protein groups were identified from individual 293T and HeLa cells on timsTOF SCP. When applied to single cholangiocarcinoma (CCA) cells and matched paracancerous cells dissociated from human fresh-frozen CCA tissue, approximately 2,000 protein groups were quantified per cell, revealing distinct metabolic and translational regulation patterns consistent with previously reported molecular features of CCA subtypes. To achieve high-throughput scProteomics, we then established an nPOP-based IBT16-TMT16 quantitative hyperplexing workflow. Across four human cell lines (293T, HeLa, A549, and LM3), our quantitative hyperplexing strategy achieved over 95% labelling efficiency and consistently identified 1,400 to 2,000 protein groups from single cells with a dynamic range spanning five orders of magnitude. In comparison to conventional multiplexing methods, our hyperplexing strategy not only enhanced proteome depth but also achieved ultrahigh-throughput (∼ 2,000 single cells per day if use Orbitrap Astral Zoom or timsUltra AIP etc.). Overall, our label-free and quantitative hyperplexing workflows provide an efficient and scalable platform for large scale scProteomics studies and clinical applications.

## INTRODUCTION

The proteome represents the dynamic molecular machinery of cells, and accurate protein characterization is crucial for understanding cellular function and regulation.^1,2^ While bulk proteomics has provided valuable insights into protein networks, it masks cell-to-cell variability by averaging signals from large populations.^3,4^ Single-cell proteomics (scProteomics) overcomes this challenge by measuring protein expression at the level of individual cells, offering new opportunities to study cellular diversity, lineage trajectories, and disease mechanisms.^5,6^ Recent advances in mass spectrometry have greatly improved the sensitivity in scProteomics, enabling label-free identification of over 5,000 proteins from single mammalian cells.^7^ However, label-free methods remain low in throughput, limiting reproducibility and scalability for large cohorts.^8,9^

To increase throughput and maintain quantitative consistency in MS detection, isobaric labelling methods such as Tandem Mass Tags(TMT),^10,11^ Isobaric Tags(IBT),^12,13^ and Isobaric Tag for Relative Absolute Quantitation(iTRAQ) ^14,15^ have been widely adopted. These multiplexed approaches encode sample-specific reporter ions in the MS2 spectra, allowing simultaneous quantification of multiple samples in a single LC–MS run.^16^ The introduction of TMTpro 18-plex has further expanded multiplexing capacity,^17,18^ and the recently released TMT 32-plex^19^ further enhances analytical throughput. However, achieving deep proteome coverage with TMT 32-plex typically requires ultrahigh-resolution mass spectrometers, which limits its broader applications in large-scale studies. Moreover, the finite number of available channels in a single isobaric reagent continues to constrain high-throughput proteomic applications. To address this, hyperplexing strategies have been developed by combining different isotopic tags or integrating multiple chemistries.^20,21^ For example, combining SILAC with TMT6 enabled the simultaneous analysis of 18 samples.^20^ Wu et al. developed a tribrid hyperplexing approach that combined TMT11, TMTpro18, and IBT16, expanding to 45 channels.^21^ More recently, integrating TAG-TMTpro with TAG-IBT16 enabled up to 102-plex analyses.^22^ However, these advances were demonstrated only in bulk proteomics and relied on extensive offline fractionation. In scProteomics, the hyperplexing for SCP (HyperSCP) workflow extended multiplexing to 28 samples by integrating stable isotope labelling by amino acids in cell culture (SILAC) with TMTpro.^23^ These advances highlight the remarkable potential of hyperplexing to overcome channel limitations and enable large-scale, quantitative scProteomics analysis.

Despite the above advances, sample preparation in scProteomics still face major challenges in handling picoliter-to-nanoliter volumes, avoiding transfer losses, and reducing reagent costs. To overcome these limitations, digital microfluidic (DMF) and droplet-based platforms have been developed.^24–26^ Early systems such as nanoliter-scale oil-air-droplet (OAD) chip^27^ and nanodroplet processing in one-pot for trace samples (nanoPOTS)^28^ improved protein recovery and proteome coverage by processing single cells in nanowells or droplets. More recently, nano-proteomic sample preparation (nPOP) has enabled parallel lysis, digestion, and labelling of thousands of single cells in picoliter droplets, reducing batch effects and reagent use.^29^ Furthermore, automated dispensing systems such as cellenONE allow the precise and reproducible manipulation of picoliter-to-nanoliter volumes. When integrated with workflows such as proteoCHIP or nPOP, they enable scalable and consistent single-cell TMT sample preparation, improving data reproducibility.^26,29^ However, the combined use of nPOP-based workflows with hyperplexing methods, which represents a promising solution for large-scale studies, has not yet been reported in scProteomics.

In this study, we established a high-throughput scProteomics workflow by integrating IBT 16 and TMTpro 16 hyperplexing with a modified nPOP method. The workflow builds on our recently reported Integral-Hyperplex method, which integrates IBT16 and TMTpro16 labelling to increase throughput by ∼30-fold and ensures accurate cross-tag quantification via normalization channels and protein-specific conversion factors.^30^ Our approach effectively expands multiplexing capacity and improves quantitative consistency. This integrated approach effectively minimizes sample transfer losses and reagent consumption while maintaining high labelling efficiency, thereby enabling sensitive and reproducible protein identification from individual cells. To assess the quantitative performance of this workflow, we profiled 480 single cells derived from four distinct cell lines (293T, LM3, HeLa, and A549), achieving deep and accurate proteome profiling.

## EXPERIMENT SECTION

### Sample Preparation of Cholangiocarcinoma (CCA) Cells

CCA samples were obtained from patients undergoing radical resection and standard lymphadenectomy at Sun Yat-sen Memorial Hospital (SYSMH). All diagnoses were rigorously confirmed through histopathological evaluation by board-certified pathologists. The use of human tissues in this study was approved by the SYSMH Ethics Review Committee (SYSKY-2023-951-01), with the requirement for written informed consent waived due to the retrospective nature of the study. Fresh-frozen CCA tissue collected from a patient were processed using the TissueGrinder system^31^ to generate enzyme-free single-cell suspensions. Following dissociation, the resulting cells were washed three times with 1× PBS to remove residual debris and contaminants.

### Proteomics Sample Preparation

For label-free scProteomics, individual clean cells were sorted into 3 μL of master mix pre-dispensed at the bottom of 384-well plates in a cellenONE system. The master mix contained 0.2% n-dodecyl-β-D-maltoside (DDM), 100 mM 4-(2-Hydroxyethyl)-1-piperazineethanesulfonic acid (HEPES), and 20 ng/μL sequencing-grade modified trypsin (Promega). The plates were subsequently sealed with self-adhesive films and incubated at 37°C with 85% humidity for 2 hours, after which the samples were stored at-80°C until LC-MS/MS analysis.

Multiplexed scProteomic samples were prepared using the nPOP workflow with modifications. Individual cells were automatically dispensed into nanoliter-scale droplets on a fluorocarbon-coated glass slide. Each droplet contained 13 nL of lysis buffer. The droplets were incubated at room temperature for 4 h to achieve efficient cell lysis and proteolytic digestion. Following digestion, 20 nL of TMT or IBT labelling reagent was added to each droplet for isobaric labelling. The labelling reaction was carried out at room temperature for 1.5 h. Subsequently, 20 nL of 5% hydroxylamine was added to each droplet to quench the reaction. After quenching, TMT and IBT labelled droplets from the same experimental group were mixed and transferred into a 384-well plate for subsequent pooling and LC–MS/MS analysis.

All multiplexed samples were combined, lyophilized under vacuum, and stored at −80 °C until further analysis. Subsequently, the samples were reconstituted in 4 µL of 0.1% (v/v) formic acid (FA) prior to LC-MS analysis.

### Data Analysis

Raw MS data were analyzed using PEAKS Online against the human UniProt FASTA database (20,375 entries). Precursor and fragment mass tolerances were set to 15 ppm and 0.05 Da, respectively. Trypsin/P was specified as the digestion enzyme, allowing up to two missed cleavages. Protein quantification was carried out using the TMT/iTRAQ Label Quantification module in PEAKS Q with de novo–assisted processing. Methionine oxidation and protein N-terminal acetylation were considered variable modifications, while TMT16 or IBT16 labels were defined as fixed modifications for the respective datasets. Labelling efficiency was evaluated using the PEAKS DB module, in which methionine oxidation, N-terminal acetylation, and TMT16 or IBT16 modifications were treated as variable. For single-cell analyses, proteins detected in fewer than 30% of normalization channels were excluded. Protein intensities were normalized to the designated channels, and missing values were imputed using a k-nearest neighbors (KNN) method.

## RESULTS AND DISCUSSION

### Optimization of LC–MS Configurations for scProteomics

We first compared the performance of timsTOF SCP and timsTOF Pro with 0.2 ng and 1 ng of HeLa cell digest, which approximate the protein content of a single somatic cell (Figure 1a). Across both input levels, the timsTOF SCP consistently achieved superior proteome coverage, identifying 3.35-fold and 1.95-fold more protein groups than the timsTOF Pro, corresponding to 542 vs 162 protein groups for 0.2 ng and 1,977 vs 1,015 protein groups for 1 ng, respectively. Next, we evaluated the effect of incorporating a trap column into the liquid chromatography setup (Figure 1b). Under the single-column configuration, markedly higher proteome identifications were obtained, with increases of 81.3% and 34.5% for the 0.2 ng (542 vs 299) and 1 ng (1,977 vs 1,470) samples Furthermore, the performance gap widened as the sample input decreased, highlighting the importance of trap-free configurations for ultralow-input proteomics, particularly at the single-cell level. In addition to sensitivity, chromatographic performance also remarkably improved under the single-column mode, as evidenced by narrower peak widths and enhanced separation resolution compared to the trap column configuration (Figure S1).

**Figure 1.**
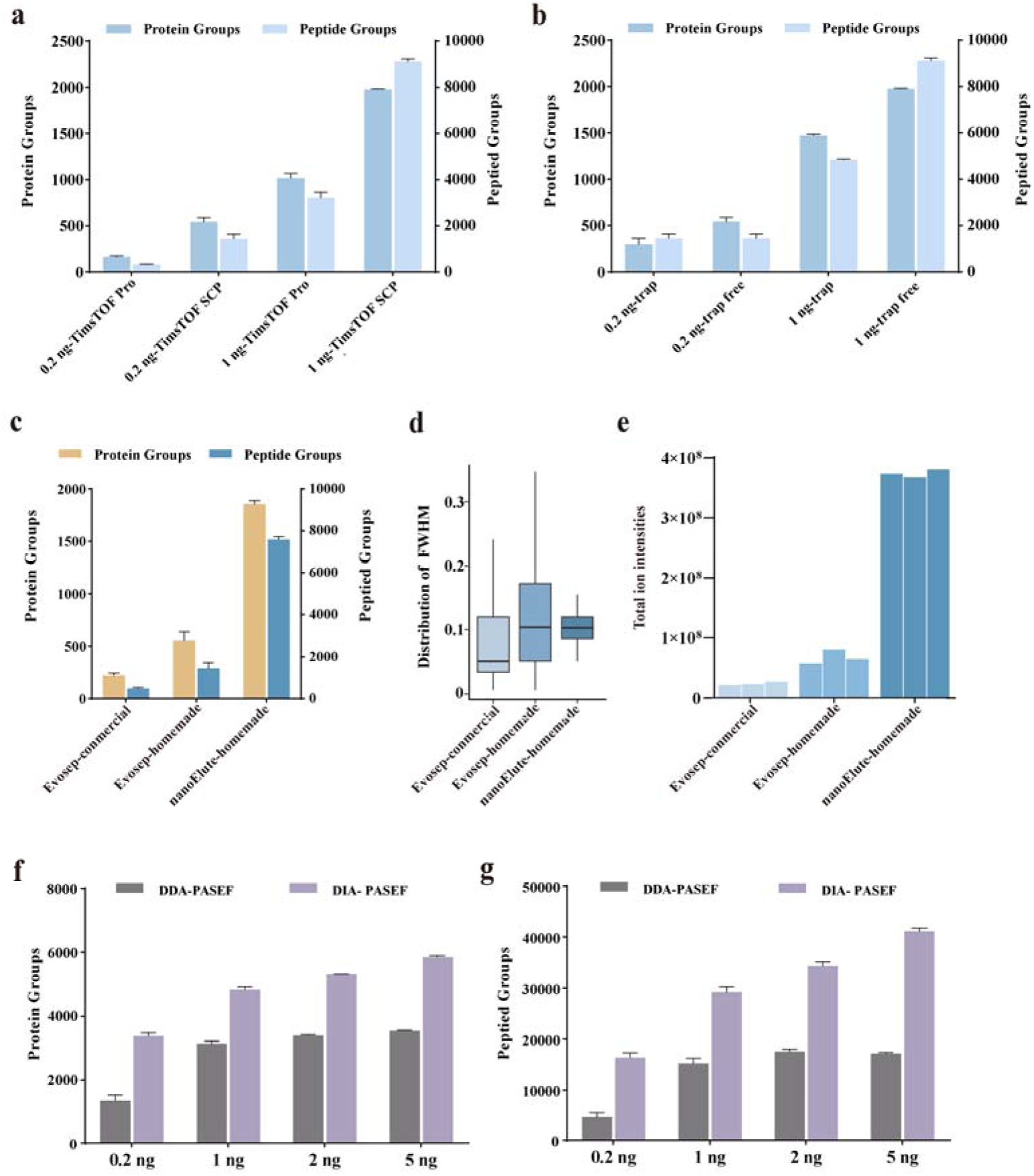
Evaluation of LC–MS strategies at the single-cell level. (a) Numbers of identified protein groups and peptide groups from 0.2 ng and 1 ng HeLa digest (n = 3), analyzed using timsTOF Pro and timsTOF SCP instruments. (b) Numbers of identified protein groups and peptide groups from 0.2 ng and 1 ng HeLa digest (n = 3), acquired using LC systems with or without a trap column. (c) Numbers of identified protein groups and peptide groups from 0.8 ng HeLa cell lysates (n = 3), analyzed on nanoElute and Evosep instruments using either commercial or homemade columns. (d) Full width at half maximum (FWHM) distributions under three distinct LC conditions. (e) Total ion intensities obtained under three distinct LC conditions. (f) Numbers of identified protein groups from 0.2 ng, 1 ng, 2 ng, and 5 ng HeLa digest, analyzed using data-dependent acquisition (DDA) and data-independent acquisition (DIA) methods. (g) Numbers of identified peptide groups from 0.2 ng, 1 ng, 2 ng, and 5 ng HeLa digest, analyzed using DDA and DIA methods.

We further compared the performance of the nanoElute and Evosep One LC systems. Each system was configured with integrated spraytip analytical columns, including either a commercial analytical column (25 cm × 75 μm i.d.) packed with 1.6 μm / 120 Å ReproSil-Pur C18 resin (Dr. Maisch GmbH, Germany) and equipped with a 10 μm Bruker ZDV integrated sprayer, or a homemade analytical column (20 cm × 50 μm i.d.) packed with 1.9 μm / 120 Å ReproSil-Pur C18 resin and equipped with a 5 μm Bruker ZDV integrated sprayer. Importantly, the nanoElute-homemade column configuration consistently delivered the best proteome coverage, achieving 1,856 protein groups and 7,850 peptide groups from as little as 0.8 ng of HeLa digest (Figure 1c). In terms of chromatographic performance, the nanoElute system exhibited narrower peak widths and distribution range, indicating higher separation efficiency (Figure 1d). In addition, we compared the total ion intensity delivered by each LC–column configuration (Figure 1e). The nanoElute–homemade column configuration generated the strongest overall ion signals, substantially exceeding both Evosep configurations. In contrast, the Evosep system produced markedly lower total ion intensity, indicating reduced ion transmission efficiency. These results further demonstrate that the nanoElute–homemade configuration offers superior signal delivery and enhanced analytical sensitivity.

Finally, we evaluated the sensitivity of different data acquisition modes across 0.2, 1, 2, and 5 ng of HeLa digest. The results demonstrated that DIA-PASEF consistently outperformed DDA-PASEF, identifying substantially higher numbers of both protein and peptide groups across all tested sample amounts (Figures 1f and g). Overall, integrating the timsTOF SCP with the nanoElute LC system using a homemade analytical column, a trap-free configuration, and DIA-PASEF acquisition resulted in superior sensitivity for scProteomic analyses.

### Deep and Reproducible Label-free scProteomics Profiling

To evaluate the performance of the cellenONE-based scProteomics workflow, we analyzed single 293T and HeLa cells. On average, more than 3,200 protein groups and over 18,000 peptide groups were identified per individual 293T cell (Figure 2a), while single HeLa cells yielded comparable depth, with approximately 3,000 protein groups and ∼15,000 peptide groups identified (Figure 2b). These results demonstrate that the label-free workflow enables highly sensitive and robust proteome coverage at the single-cell level. To further assess data quality, we performed principal component analysis (PCA) across all single-cell and bulk samples. As shown in Figure 2c, the three sample types (bulk HeLa digests, single 293T cells, and single HeLa cells) were accurately clustered and clearly separated, indicating distinct proteomic profiles. The coefficient of variation of the three groups all fell within a relatively low range (Figure S2). Consistently, correlation analysis revealed high intra-group reproducibility, with correlation coefficients close to 1 within the same cell type, whereas inter-group correlations were markedly lower (Figure 2d). In addition, unsupervised hierarchical clustering based on quantified proteins accurately grouped the samples according to their biological origin (Figure 2e). In summary, these results demonstrate that the cellenONE-based scProteomics workflow provides deep proteome coverage, excellent quantitative reproducibility, and clear biological discrimination across different cell types. The routine identification of over 3,000 protein groups per single mammalian cell on the timsTOF SCP highlights the high sensitivity and robustness of our workflow.

**Figure 2.**
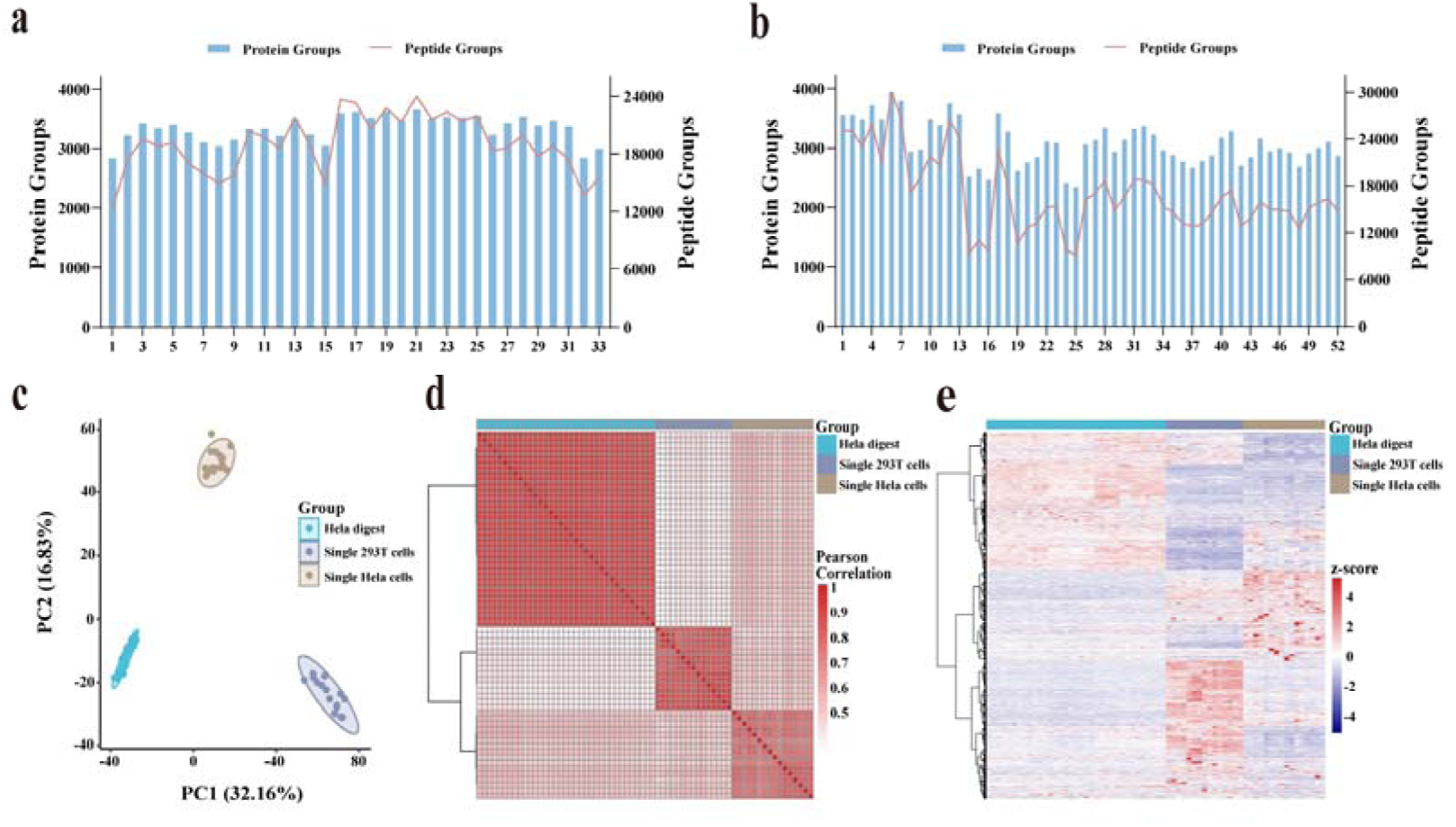
Performance of Label-free scProteomics. (a–b) Numbers of identified protein groups (blue bars) and peptide groups (red lines) from single 293T (a) and HeLa (b) cells. (c) PCA analysis of scProteomic from 293T cells, HeLa cells, and bulk HeLa digest. (d) Pearson correlation analysis among the three samples. (e) Hierarchical clustering heatmap of quantified proteins across the three samples.

### Label-free scProteomics to Examine Proteome Diversity in Cholangiocarcinoma

Cholangiocarcinoma (CCA) is the second most prevalent primary malignant tumor of the liver, characterized by pronounced heterogeneity, aggressive invasiveness, and poor prognosis.^32–36^ Within CCA, the tumor microenvironment exhibits significant heterogeneity among distinct cell populations within the same tumor. Such heterogeneity results in uneven immune cell infiltration, variable stromal cell composition, and heterogeneous distributions of stem-like tumor cells, ultimately presenting substantial challenges for the development of effective therapeutic strategies.^37^

To characterize proteomic alterations in distinct cell types within the tissue microenvironment at the single-cell level, we applied a cellenONE-based scProteomics workflow to analyze individual CCA cells together with their matched paracancerous counterparts. The scProteomic profiling of CCA and matched paracancerous cells achieved the identification of approximately 1,000–2,800 protein groups and 5,000–18,000 peptide groups (Figure 3a), while revealing pronounced cell-to-cell variability. PCA revealed clear separation between cancer derived and paracancer cells (Figure 3b). Moreover, hierarchical clustering and correlation analyses separated the two sample classes while also resolving intratumoral subclusters (Figure 3c and 3d). Given the pronounced intratumoral heterogeneity of CCA and the need to decode cell-type-specific proteomic alterations within its microenvironment, we further performed differential expression and functional enrichment analyses on scProteomic data. As shown in Figure 3e, numerous proteins exhibited significant differential expression between the two groups. Red dots represent upregulated proteins (e.g., HSPD1 and PDHA1) implicated in cellular stress responses and energy metabolism, processes closely associated with the aggressive invasiveness of CCA. Blue dots represent downregulated proteins (e.g., CYP4F3 and CES1), key enzymes involved in drug metabolism whose dysregulation may contribute to the poor therapeutic response observed in CCA. Gene Ontology (GO) enrichment analysis (Figure 3f) further revealed that these upregulated proteins were significantly enriched in pathways critical for CCA progression, including biological oxidation, fatty acid metabolism, steroid metabolism, and eukaryotic translation initiation and elongation. These pathways likely support the malignant phenotypes of CCA cells and the maintenance of intra-tumoral subpopulations. In contrast, downregulated pathways were less prominent.

**Figure 3.**
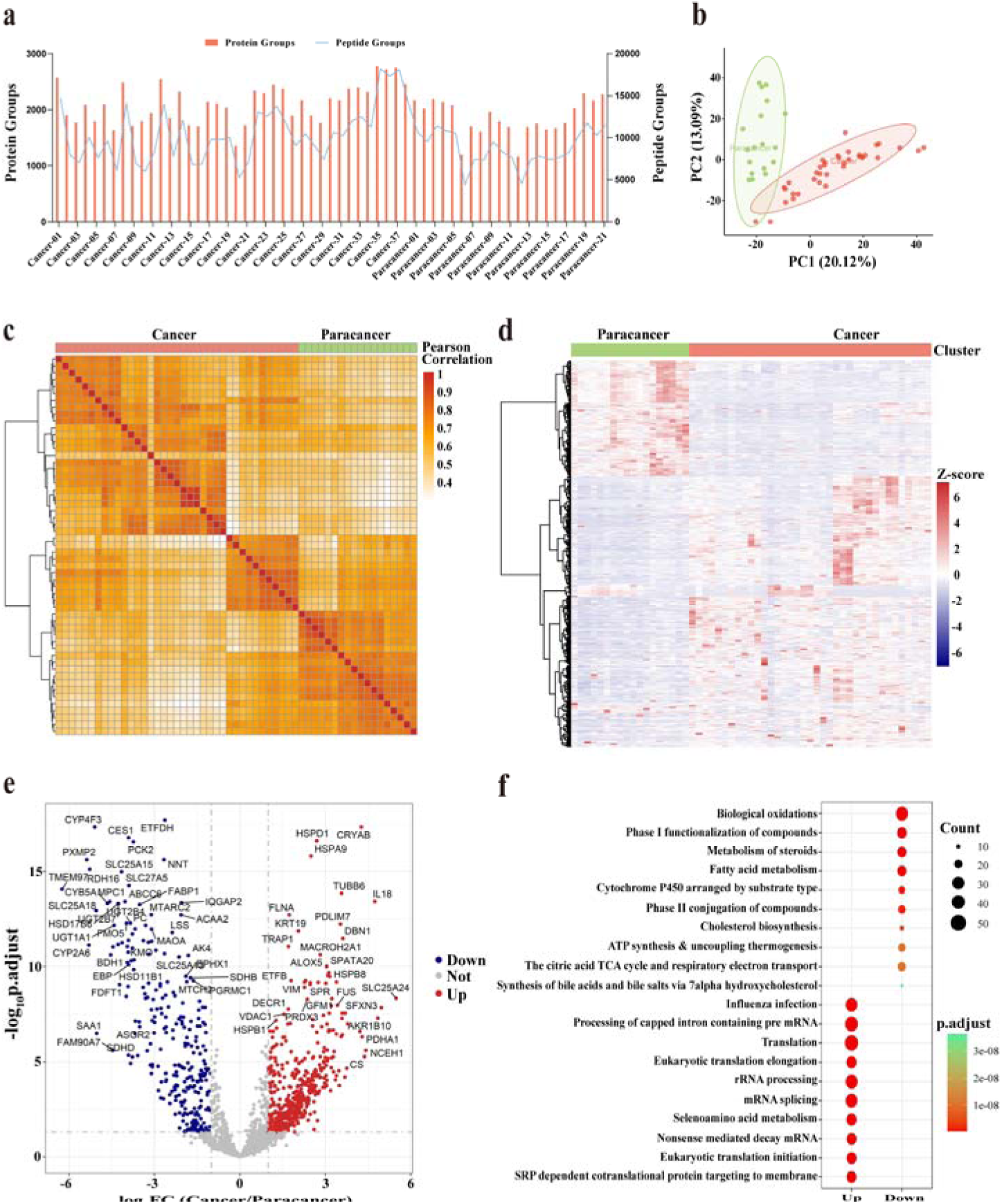
Proteomic profiling of single cholangiocarcinoma and paracancer cells. (a) Numbers of identified protein groups and peptide groups from single cholangiocarcinoma and paracancer cells. (b) PCA of the two cell types. (c) Pearson correlation analysis among the two cell types (d) Hierarchical clustering heatmap of quantified proteins across the two cell types. (e)Volcano plot showing differentially expressed proteins CCA samples compared paracancer samples. (f) GO enrichment analysis of biological processes in CCA samples compared to paracancer sample.

Our results are consistent with recent single-cell studies of CCA. Previous single-cell analyses have reported pronounced intratumoral heterogeneity in CCA, including distinct malignant subpopulations with divergent metabolic and stress-response programs^38,39^. In line with these observations, our scProteomic analysis revealed clear separation between cancer and paracancerous cells, as well as evident intratumoral subclusters, accompanied by differential regulation of metabolic, mitochondrial, and protein homeostasis–related pathways.^35,40^ Together, these results support the notion that proteomic heterogeneity and metabolic reprogramming are prominent features of CCA at the single-cell level.

Overall, these results indicate that CCA progression is accompanied by extensive proteomic reprogramming, characterized by enhanced proteostasis capacity, cytoskeletal reorganization, and altered energy metabolism, coupled with pronounced heterogeneity at single-cell resolution. Such features are consistent with increased phenotypic plasticity and functional diversity among tumor cells. Importantly, this study highlights the unique strength of scProteomics in uncovering cell-type-specific proteomic features, such as dependence on chaperone networks, metabolic nodes, or stress-response pathways, that are frequently masked in bulk proteomics analyses.

### Integration of nPOP workflow with Quantitative Hyperplexing for High-throughput scProteomics Analysis

Though the above label-free workflow achieved high sensitivity and high simplicity, it is not feasible for high-throughput large-scale scProteomics applications. To obtain high-throughput, we integrated nPOP method with IBT16-TMTpro16 hyperplexing (Figure 4a), which can further expand multiplexing depth in scProteomics while preserving quantitative accuracy. This strategy enables precise single-cell isolation, nanoliter-scale reactions, and efficient peptide labelling with minimal sample loss. Due to the resolution limitation of the timsTOF SCP, the two 16-plex reagents were divided into two groups to avoid channel overlap during MS acquisition. For IBT16, Group A consisted of channels 114 (reference), 115C, 116C, 117C, 118C, 119C, 120C (blank), and 121C (carrier), whereas Group B included 115N (carrier), 116N, 117N, 118N, 119N, 120N, 121N (blank), and 122 (carrier). For TMTpro16, Group A included 126 (reference), 127C, 128C, 129C, 130C, 131C, 132C (blank), and 133 (carrier), while Group B contained 127N (reference), 128N, 129N, 130N, 131N, 132N, 133N (blank), and 134N (carrier).

**Figure 4.**
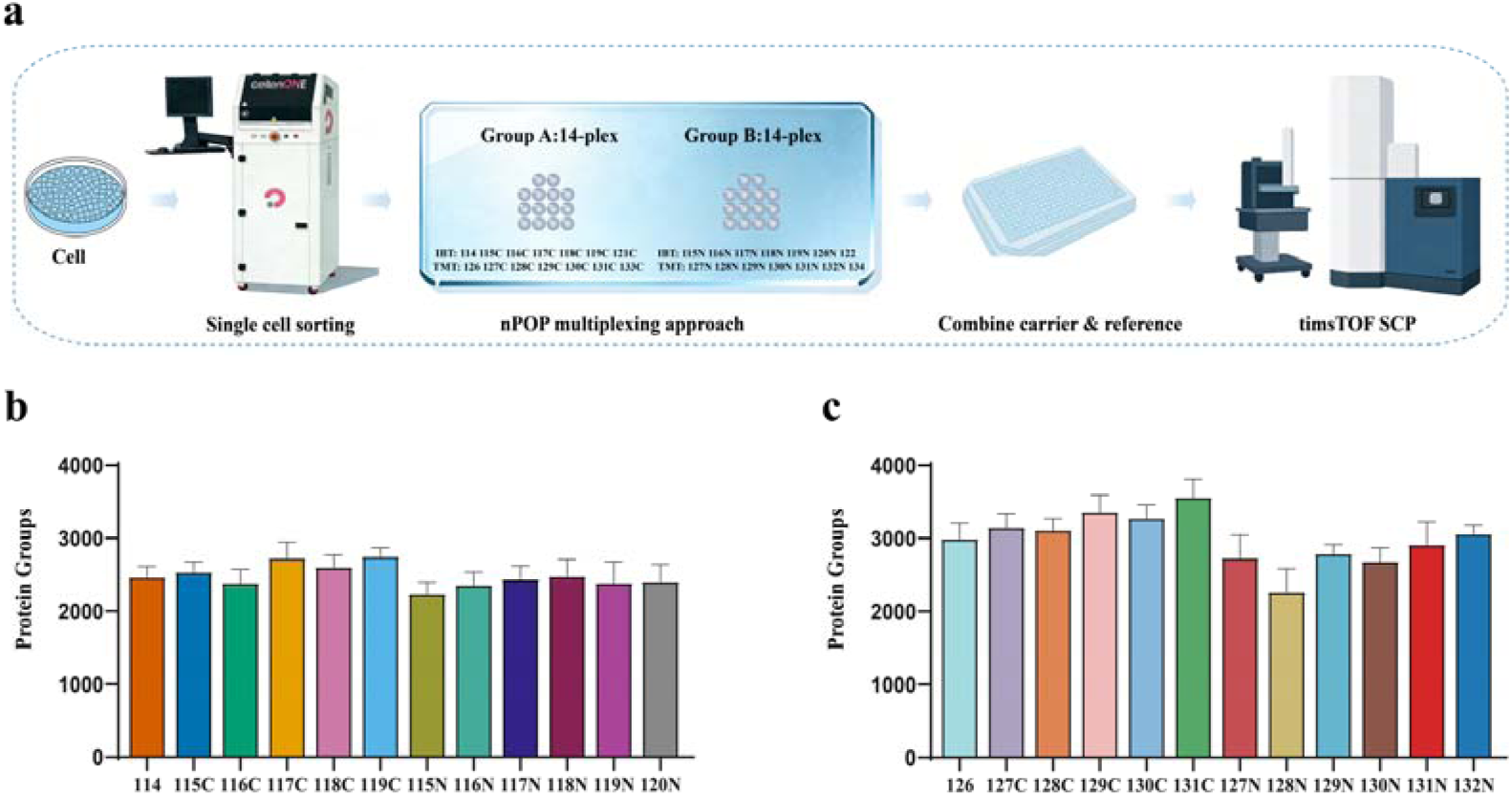
Schematic illustration of hyperplexing scProteomic analysis using the nPOP-based labelling strategy. (a) Workflow of the nPOP-based labelling strategy. The workflow includes single-cell isolation, lysis, digestion, labelling, and proteomic analysis. (b–c) Number of identified protein groups per single 293T cell labelled with IBT16 (b) or TMTpro16 (c).

To systematically evaluate the quantitative performance of each individual regent labelling, single 293T cells were analyzed using the nPOP-based labelling workflow on the CellenONE platform. As shown in Figures 4b and 4c, both labelling approaches exhibited stable protein identification across channels, with ∼3,000 protein groups quantified per channel. Notably, the TMTpro16-labelled samples showed slightly deeper proteome coverage than the IBT16-labelled counterparts, indicating improved ion response under identical LC–MS conditions. Both labelling strategies achieved high and consistent labelling efficiencies (Figure S3), demonstrating their robustness for scProteomics analysis. The proteome coverage obtained in this study notably exceeds that of the recently developed high-throughput nPOP workflow, which identified approximately 1,000 proteins per human cell using TMTpro 32-plex labelling.^29^

These results demonstrate that the nPOP-based workflow supports robust and high-efficiency labelling reactions for both IBT16 and TMTpro16 reagents, achieving consistent scProteomics coverage.

### Analysis of Diverse Cell Types Using the Quantitative Hyperplexing Strategy

Next, to further evaluate the performance of the hyperplexing strategy, we applied the established workflow to multiplexed proteomic analysis of four human cell lines (LM3, HeLa, A549, and 293T) as a proof-of-concept application. As shown in Figure 5a, an average of 1,400-2,000 protein groups were quantified per cell. In addition, labelling efficiencies exceeded 95% across all four cell types (Figure S4), demonstrating that this workflow enables stable and highly efficient labelling reactions across diverse cellular contexts.

**Figure 5.**
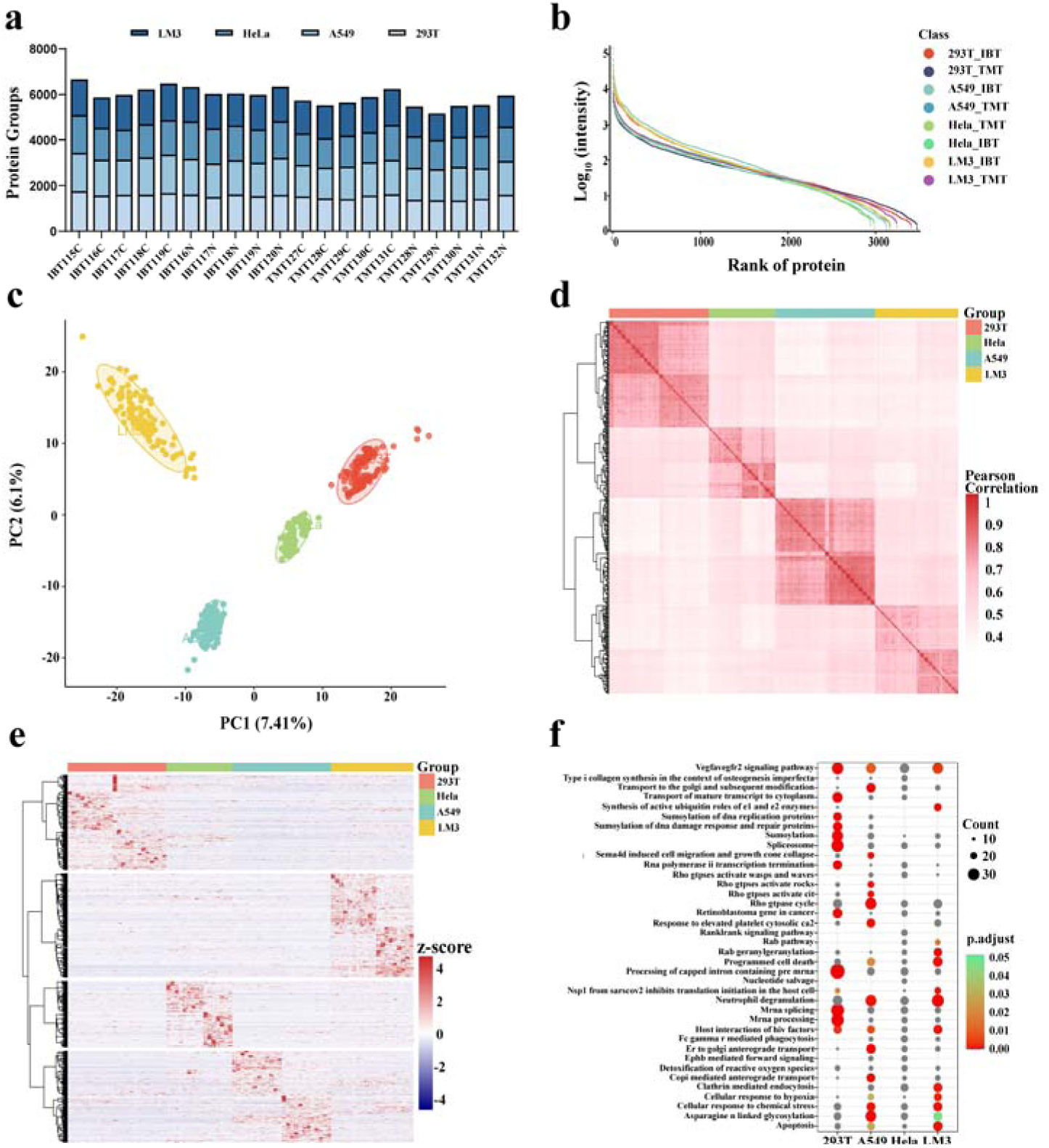
Quantitative performance of the nPOP-based hyperplexing strategy across four human cell lines. (a) Protein quantification results for the combined IBT–TMT labelling of four human cell lines (LM3, HeLa, A549, and 293T) (b) Dynamic ranges (c) PCA analysis, (d) Pearson correlation analysis among four cell types, (e-f) Unsupervised hierarchical clustering of proteins and gene ontology biological processes enrichment analysis of the protein clusters from the heatmap.

Importantly, the dynamic ranges of quantified protein groups from single cells achieved five orders of magnitude (Figure 5b), indicating that both methods effectively captured proteins across a wide range of abundances levels. PCA analysis consistently and accurately separated the four cell types according to their biological identities rather than TMT or IBT labelling, indicating that our strategy reliably distinguishes TMT and IBT derived reporter ions and effectively captures the substantial biological differences among cell types (Figure 5c). Pairwise correlation heatmaps showed high consistency within each group (Figure 5d), while hierarchical clustering revealed clear differences in proteomic profiles among the different cell lines (Figure 5e). Moreover, GO enrichment analysis revealed pronounced cell-type-specific functional signatures across the four human cell lines analyzed (Figure 5f). In 293T cells, significantly enriched terms were predominantly associated with RNA metabolism, including mRNA processing, splicing, and translation-related pathways, consistent with their high transcriptional and translational activity. HeLa cells displayed enrichment of pathways linked to genome maintenance, such as DNA replication, chromatin-associated processes, and cell cycle regulation, reflecting their highly proliferative phenotype. In contrast, A549 cells showed preferential enrichment of metabolic pathways, stress-response programs, and signaling cascades related to growth factor and hypoxia responses, in line with their lung adenocarcinoma origin and metabolic adaptability. Notably, LM3 cells were characterized by enrichment of pathways involved in extracellular matrix organization, cell migration, vesicle-mediated transport, and apoptosis-related processes, highlighting features associated with invasiveness and metastatic potential. Together, these results demonstrate that the our hyperplexing strategy effectively captures biologically meaningful proteomic heterogeneity across diverse cell states.

Overall, the IBT16-TMTpro16 hyperplexing strategy markedly enhanced both the throughput and quantitative depth of scPproteomics, expanding the multiplexing capacity to 32 channels per run and enabling more efficient parallel sample analysis. Across four human cell lines, this workflow consistently quantified 1,400-2,000 protein groups per cell with labelling efficiencies exceeding 95%, while accurately capturing proteins across a dynamic range spanning five orders of magnitude. Principal component and hierarchical clustering analyses revealed clear separation among cell types and high biological reproducibility, whereas GO enrichment analysis further confirmed the distinct biological functions of cell-type-specific proteins.

## CONCLUSION

To obtain high sensitivity in label-free scProteomics, we optimized the chromatographic and MS parameters for scProteomics analysis. Under these conditions, more than 3,000 protein groups were identified from individual 293T and HeLa cells on average. In the proteomic analysis of single CCA cells and their matched paracancerous cells, we achieved highly sensitive detection and identified approximately 2,000 protein groups per cell. Our data revealed marked differences in metabolic and translational regulation among tumor subpopulations, which are consistent with previously reported molecular features of cholangiocarcinoma subtypes.^39^

To achieve ultrahigh-throughput scProteomics, we developed an integrated IBT16-TMT16 hyperplexing strategy based on the nPOP microdroplet workflow. This approach achieved labelling efficiencies exceeding 95% across multiple human cell types and expanded the effective number of quantification channels to 32 per LC-MS run. Compared with conventional multiplexed or label-free approaches, our hyperplexing strategy significantly enhanced both proteome depth and quantitative precision, enabling consistent identification of 1,400-2,000 protein groups per cell, with a dynamic range spanning five orders of magnitude. Notably, our workflow enables the preparation of up to ∼1,440 single cells in a single batch using cellenONE, when coupled with the hyperplexing strategy, supporting an MS detection throughput of approximately 2,000 cells per day if use state-of-the-art MS instruments like Orbitrap Astral Zoom or timsUltra AIP. This quantitative hyperplexing workflow markedly improves throughput and data robustness relative to conventional multiplexed methods. Furthermore, our approach offers a scalable platform for higher multiplexing levels (e.g., TMTpro 32-plex and IBT32-plex) and compatibility with other labelling reagents, enabling large-scale biological and clinical applications.

## ASSOCIATED CONTENT

### Data availability

The mass spectrometry proteomics data related to this study have been deposited to iProX database (https://www.iprox.cn/page/PSV023.html) under the accession number IPX0015905000.

### Supporting Information

The Supporting Information is available free of charge at…

Partial Experimental section, partial results (Figures S1−S4) of method evaluation and classifier performance comparison (PDF).

### Author Contributions

Kunpei Cai: Methodology, Investigation, Formal analysis, Writing-Original Draft. Qing Zeng: Methodology, Investigation. Chuanxi Huang: Formal analysis. Jing Yang: Supervision. Fuchu He: Supervision. Yun Yang: Methodology, Conceptualization, Supervision, Writing–Review & Editing.

## ACKNOWLEDGMENTS

This work was supported by National Key R&D Program of China (2025YFA1309400), Major Project of Guangzhou National Laboratory (Grant No. GZNL2024A03001), the Ministry of Science and Technology of the People’s Republic of China (Grant No. 2020YFE0202200), the Pre-study Project of Phronesis Medicine Large-scale Scientific Facility funded by Guangzhou Development District.

## Notes

The authors declare no competing financial interest.

