## Supplementary Material for "High-throughput Single-cell Proteomics Enabled by Integrating nPOP workflow with Quantitative Hyperplexing"

**Corresponding Authors**

### Contents

**EXPERIMENTAL SECTION** **S3**

**Figure S1.** **S6**

**Figure S2.** **S7**

**Figure S3.** **S8**

**Figure S4.** **S9**

#### ■ EXPERIMENTAL SECTION

#### Materials.

TMTpro 16-plex, TMT0 labelling reagents, and 50% hydroxylamine solution were obtained from Thermo Fisher Scientific (USA). The IBT 16-plex isobaric reagents were supplied by Nanjing Apollomics Biotech Inc. (China).

##### Cell Culture and Harvest.

Human embryonic kidney (HEK) 293T cells, HeLa (human epithelial carcinoma) cells, A549 cells (human lung adenocarcinoma epithelial cell line) and LM3 cells (human hepatocellular carcinoma cell line with high metastatic potential) were maintained in Dulbecco's Modified Eagle Medium (DMEM) containing 10% fetal bovine serum (FBS) and 1% penicillin-streptomycin at 37°C with 5% $CO_2$. For single-cell isolation, cells were trypsinized, washed three times with phosphate buffered saline (PBS), and resuspended in PBS at 200 cells/μL for processing on the CellenONE system.

#### Bulk Sample Preparation for Carrier and Reference channels.

Cells were detached with 0.05% trypsin–ethylenediaminetetraacetic acid (EDTA) and washed three times with PBS. Pelleted cells were lysed in 1% Sodium deoxycholate (SDC), 100 mM tris(2-carboxyethyl)phosphine (TCEP, pH 7.0), 40 mM chloroacetamide (CAA), and 100 mM tris(hydroxymethyl)aminomethane hydrochloride (Tris-HCl) (pH 8.0), followed by sonication. Protein content was quantified using a NanoDrop One, and samples were digested overnight at 37 °C with trypsin at a 1:30 enzyme-to-protein ratio. Peptides were acidified (pH < 3) and desalted using Sep-Pak tC18 cartridges. Carrier peptides were labeled with TMTpro-133C, TMTpro-134N, IBT16-121C, or IBT16-122, and reference peptides with TMTpro-126, TMTpro-127N, IBT16-114, or IBT16-115N. All reactions were quenched with 5% hydroxylamine before pooling.

#### NanoLC-MS/MS analysis on timsTOF SCP.

NanoLC-MS/MS was performed on a timsTOF SCP (Bruker Daltonics) with a CaptiveSpray nano-ESI source. Peptides were desalted online using a 300 µm × 5 mm C18 PepMap100 trap column (Thermo Fisher Scientific). Separation was conducted on a nanoElute UHPLC system using an integrated spray-tip C18 column (50 µm × 20 cm, 1.9 µm, 120 Å; Dr. Maisch). Mobile phases were water with 0.1% formic acid (A) and acetonitrile with 0.1% formic acid (B). The flow rate was 100 nL/min, except for the first 5 min at 200 nL/min. A 60-min gradient was applied: 5–8% B (0–5 min), 8–10% (5–8 min), 10–26% (8–47.6 min), 26–45% (47.6–53 min), 45–95% (53–55 min), and 95% (55–60 min). The column temperature was 60 °C.

MS acquisition was performed in DDA mode over m/z 100–1700 with an ion mobility range of 1.3–0.7 Vs $cm^{-2}$. Each cycle comprised one TIMS-MS scan and ten PASEF MS/MS scans with a 166 ms ramp. Precursors with charge states 0 to +5 were selected. Collision energy increased with mobility from 44.8 eV at $1/K_0$ = 0.60 Vs $cm^{-2}$ to 89.6 eV at $1/K_0$ = 1.60 Vs $cm^{-2}$.

#### Data analysis

DIA data from method development were analyzed using Spectronaut 18 (Biognosys) software. These data were searched against the human UniProt FASTA database (20,375 entries, downloaded on December 3, 2021). Cysteine carbamidomethylation was set as fixed modification. N-terminal acetylation of protein and methionine oxidation were set as variable modifications. Two missed cleavages were allowed with trypsin/P digestion. Precursor filtering was set to perform based on Q-values. Quantification results were output with protein and peptide false discovery rates (FDRs) less than 0.01. All other settings were left at their default values.

DDA data from label-free samples were processed using MaxQuant (version 2.0.3.0) with the integrated Andromeda search engine. Spectra were searched against the

UniProt Homo sapiens reference proteome database (Swiss-Prot, 20,375 reviewed protein entries, release 2021_03). Trypsin/P was specified as the proteolytic enzyme allowing up to two missed cleavages. Carbamidomethylation of cysteine was set as a fixed modification, while methionine oxidation and protein N-terminal acetylation were defined as variable modifications.

#### Bioinformatics and statistical analysis

Proteins identified in fewer than 50% of samples were removed prior to downstream analyses. Protein abundance values were median-normalized across samples, and missing values were imputed using the k-nearest neighbours (KNN) algorithm. The resulting quantitative matrix was subsequently $log_2$-transformed.Differential expression analysis was conducted using two-tailed unpaired Student's t-tests or one-way ANOVA, as appropriate. Proteins with a fold change > 1.5 and $p < 0.01$ were considered differentially expressed proteins (DEPs); for ANOVA, significance was defined as a false discovery rate–adjusted q-value < 0.05. All statistical analyses and data visualization were performed in R (v4.2.2). The following R packages were used: pheatmap (v1.0.12), ggplot2 (v3.4.4), pROC (v1.18.5), and clusterProfiler (v4.2.2). Gene Ontology (GO) enrichment analysis was conducted using the msigdbr (v7.5.1) and clusterProfiler packages, with a significance threshold of $q < 0.05$. Gene expression clustering was performed using Mfuzz (v2.54.0). Statistical analyses and graphical representation of the five reported biomarkers were generated using GraphPad Prism (v8.0).

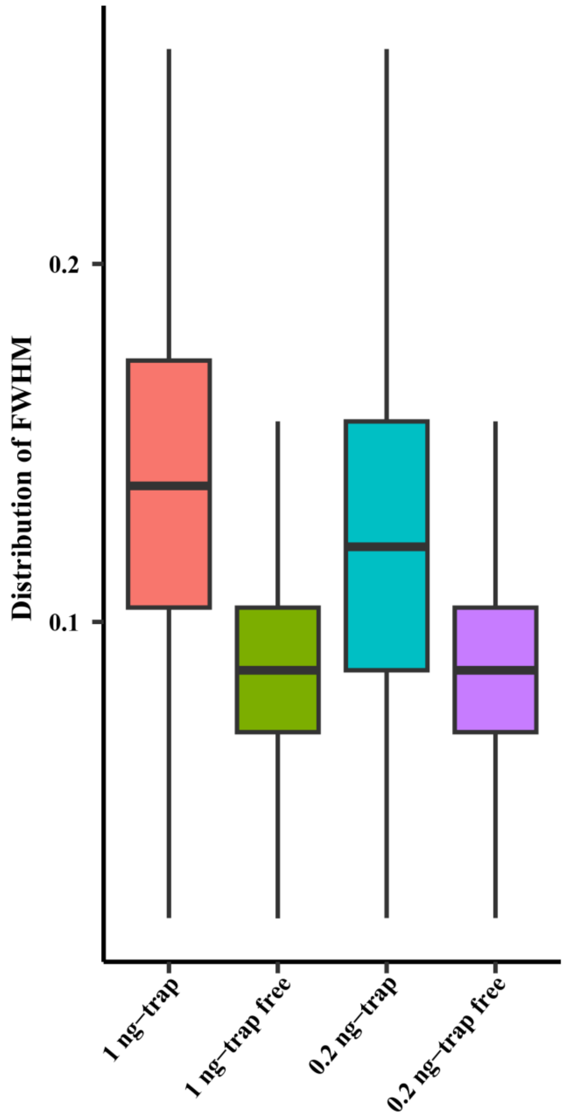


**Figure S1.** Comparison of the distribution of Full Width at Half Maximum (FWHM) between LC configurations with and without an external trap column.

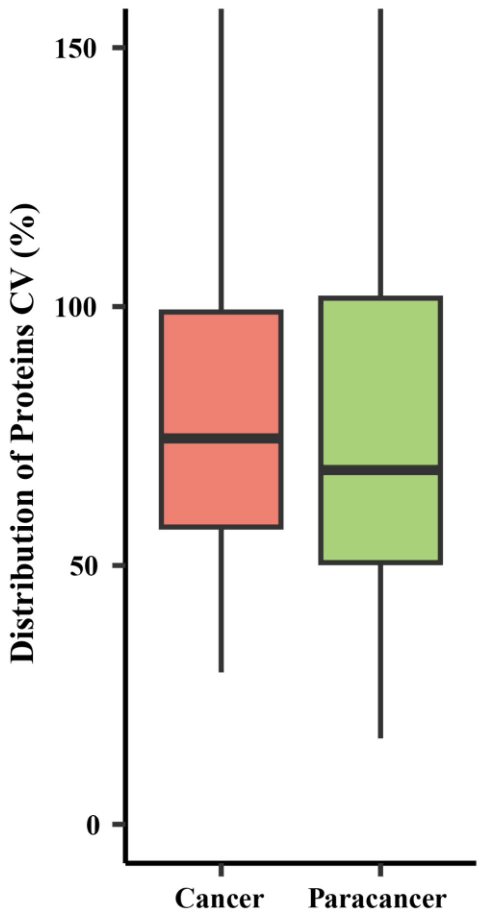


**Figure S2.** Distribution of protein coefficient of variation (CV, %) in single cells from cancer and paracancer tissues.

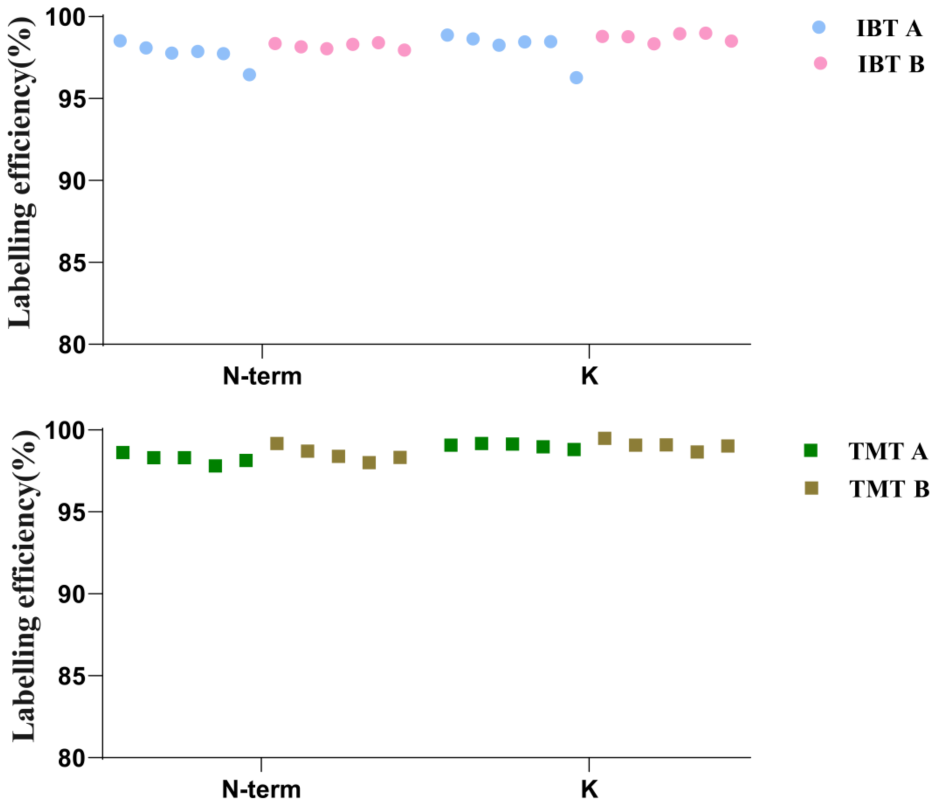


**Figure S3.** Labelling efficiencies at peptide N-termini and lysine residues for single 293T cells processed using the nPOP workflow with different labelling reagents. (a) IBT 16 labelling. (b) TMTpro 16 labelling.

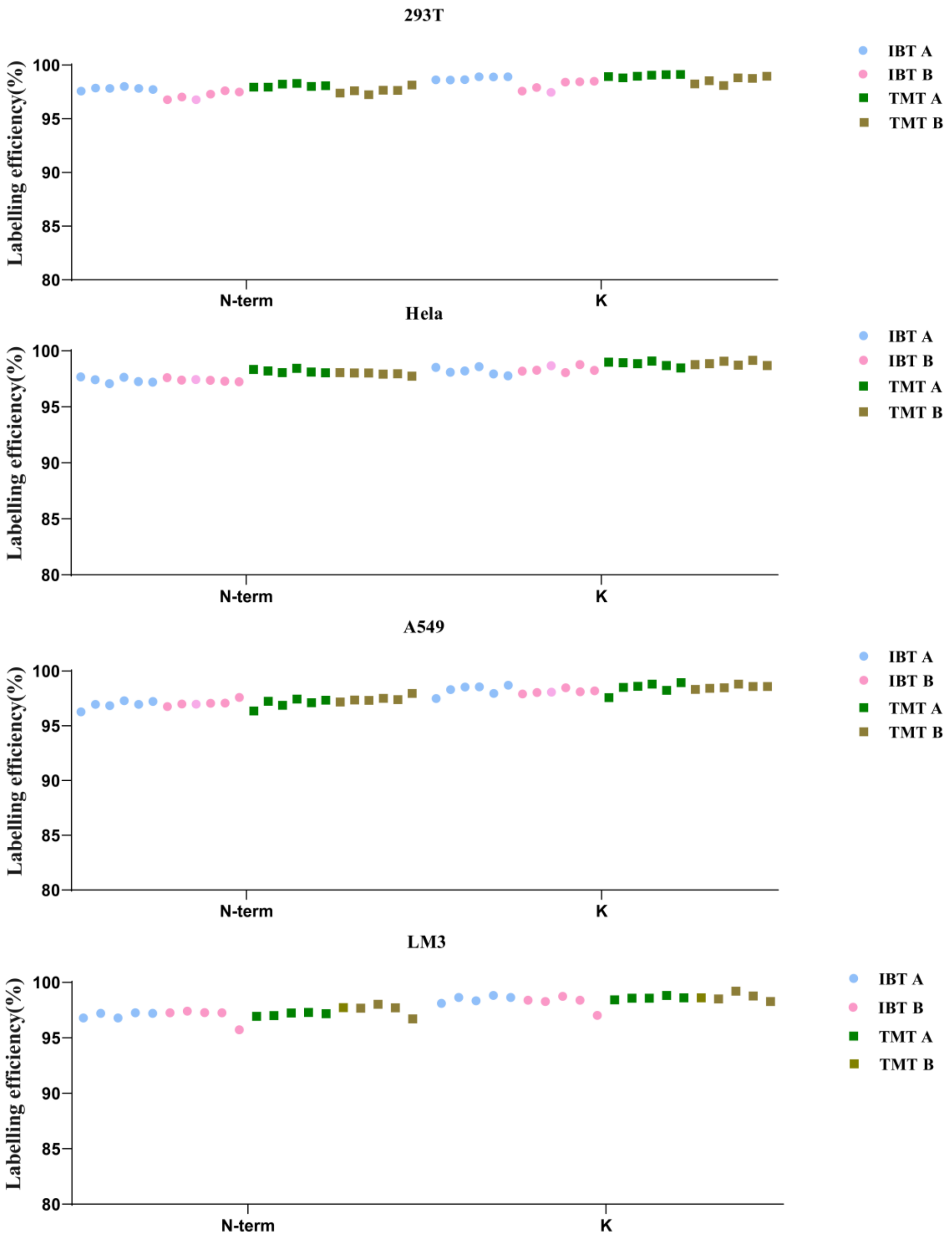


**Figure S4.** Labelling efficiencies of single cells processed and labelled based on the nPOP workflow. Labelling efficiencies on peptide N-termini and lysine residues for (a) 293T, (b) HeLa, (c) A549 cells, and (d) LM3 cells. Circles indicate IBT labelling, and squares indicate TMT labelling.
